# HOXB13 alters chromatin accessibility in prostate cancer through interactions with the SWI/SNF complex

**DOI:** 10.1101/2023.09.04.556101

**Authors:** Shreyas Lingadahalli, Betul Ersoy Fazlioglu, Umut Berkay Altintas, Ahmet Cingoz, Emirhan Tekoglu, Ivan Pak Lok Yu, Ugur Meric Dikbas, Hans Andomat, Ibrahim Kulac, Tunc Morova, Kevin Xiao, Martin Gleave, Ladan Fazli, Paloma Cejas, Artem Cherkasov, Wilbert Zwart, Henry Long, Colin Collins, Tugba Bagci-Onder, Nathan A. Lack

## Abstract

HOXB13 is a posterior homeobox protein that is associated with the initiation and growth of prostate cancer (PCa). While most research has focused on the role of HOXB13 on androgen receptor (AR) activity, we demonstrate that HOXB13 is essential to the proliferation of both AR-positive and -negative PCa. Strikingly, HOXB13 is remarkably selective and has almost no effect on non-prostatic tissues. Despite this common essentiality in PCa, HOXB13 activity is markedly different in AR-negative PCa, where interactions with the AP-1 change the HOXB13 cistrome in stem-cell like castration-resistant prostate cancer. We show that HOXB13 activity is commonly mediated by SMARCD2, a member of the mSWI/SNF chromatin remodeling complex. Despite the distinct transcription factor interactions in AR-positive and -negative PCa the HOXB13/SMARCD2 commonly alters chromatin accessibility at HOXB13 binding sites that causes increased proliferation in PCa. Overall, this work demonstrates a novel mechanism of action for HOXB13 and highlights its critical role in AR-negative castration-resistant prostate cancer.

## INTRODUCTION

Prostate cancer (PCa) is the second leading cause of cancer-related death in North American men (1). While androgen receptor pathway inhibitors (ARPI) are the standard-of-care to treat recurrent or metastatic PCa, the clinical response is unfortunately only temporary, and the majority of patients (>80%) progress to a lethal castration-resistant PCa (CRPC) (2,3). There is an unmet need for new therapeutic targets to treat aggressive PCa that are independent of androgen receptor (AR).

Prostate embryogenesis is mediated through the coordinated expression of posterior homeotic (HOX) genes including *HOXD13*, *HOXC13*, and *HOXB13* (4–6). Unlike most HOX genes that are expressed during development, *HOXB13* remains highly transcribed in the adult prostate where it is believed to play a role in cell identity (4). There is extensive clinical and experimental evidence that HOXB13 contributes to PCa development. Several *HOXB13* germline variants, including G84E, have been associated with increased PCa frequency, Gleason scores, and a greater likelihood of positive surgical margins at the time of radical prostatectomy (7–10). HOXB13 is proposed to contribute to neoplastic transformation through alterations in coregulatory protein interactions. During PCa progression, HOXB13 transitions from interacting with the tumour suppressor three-amino-acid loop extension protein, MEIS1, to the oncogenic AR (11–13). The AR cistrome is dramatically altered through these interactions leading to locus-specific changes in AR-mediated transcription that contribute to PCa development and growth (14,15). HOXB13 also interacts with a constitutively active AR splice variant and alters both DNA binding and transcription of AR-V7 target oncogenes (16). While there is a clear relationship between HOXB13 and AR there are conflicting studies if *HOXB13* is itself regulated by AR (17–19). Further, recent work demonstrated that HOXB13 also has an AR-independent function that modulates lipogenesis in PCa (20). Despite the critical role of this transcription factor, there is controversy about the role of HOXB13 in PCa with some studies showing that HOXB13 promotes PCa growth (14,15,21), while others suggesting it suppresses AR signaling and tumor growth (11,20,22).

In this study, we demonstrate that HOXB13 is critical to the growth and proliferation of both AR-dependent and -independent PCa models. We show that HOXB13 activity in these models dramatically changes through interactions with cell-specific transcription factors. Yet despite the diverse transcription factor interactions, HOXB13 activity is commonly mediated by SMARCD2, a member of the mSWI/SNF chromatin remodeling complex. This HOXB13/SMARCD2 interaction is stabilized through interactions with cell-specific transcription factors which alters chromatin accessibility at HOXB13 binding sites that causes increased proliferation in PCa.

## MATERIAL AND METHODS

### Cell culture

LNCaP, PC3, 22Rv1, VCaP, DU145, HEK293T, and Hep3B cell lines were purchased from ATCC and validated by short tandem repeat testing. LNCaP-ABL (23) and LNAI (24) cell lines were a kind gift from Professor Zoran Culig and Professor Paul Rennie. LNCaP cells were cultured in RPMI 1640 with L-glutamine (Lonza, Switzerland #12-702Q) supplemented with 10% tetracycline-free fetal bovine serum (Biowest, Franse #S181T) and 1% penicillin/streptomycin (Gibco, USA #15140122). PC3, 22Rv1, DU145, and Hep3B cells were cultured in RPMI 1640 with L-glutamine (Lonza, Switzerland #12-702Q) supplemented with 10% fetal bovine serum (Biowest, France #S1300) and 1% penicillin/streptomycin (Gibco, USA #15140122). VCaP and HEK293T cells were cultured in DMEM-high glucose (Sigma-Aldrich, USA #D6429) supplemented with 10% fetal bovine serum (Biowest, Franse #S1300) and 1% penicillin/streptomycin (Gibco, USA #15140122). LNCaP-ABL and LNAI cells were cultured in RPMI 1640 with L-glutamine (Lonza, Switzerland #12-702Q) supplemented with 10% charcoal-stripped fetal bovine serum (Biowest, Franse #S181F) and 1% penicillin/streptomycin (Gibco, USA #15140122). All cell lines were regularly tested for mycoplasma.

### Plasmids and cloning

pLENTI CMV V5-Luc Blast vector (Addgene #19785) and lentiCRISPRv2 (Addgene #52961) were used for CRISPR/Cas9 genome editing. lentiCRISPRv2 vector was digested with BsmBI (NEB, USA #R0580) and dephosphorylated with Antarctic Phosphatase (NEB, USA #M0289S). Annealed sgRNA oligo pairs were phosphorylated with T4 PNK (NEB, USA #M0201S) and ligated with Quick Ligase (NEB, USA #M220S). pLKO_HOXB13_#1 (Addgene #70093) and pLKO_HOXB13_#2 (Addgene #70094) were used for shRNA knock-down. For shRNA control; shFF was cloned into pLKO.1-puro (Addgene #8453). For tet-inducible shRNA knock-down system; shCtrl and shHOXB13 were cloned into Tet-pLKO-puro (Addgene #21915). sgRNA and shRNA sequences can be found in Supplementary Table 1. HOXB13 was PCR amplified from pLX302_HOXB13 (Addgene #70089), digested with SalI/XbaI and ligated into linearized pENTR1A (Addgene #17398). Site-directed mutagenesis was performed to generate mutation constructs. Entry plasmids were recombined to the all-in-one Tet-inducible pLX402 (Addgene #25896) destination vector with Gateway LR Clonase II Enzyme Mix (Invitrogen, USA #11791020). All cloning was validated by Sanger sequencing. gRNA/shRNA sequences are given in Supplementary Table 1.

### Viral packaging and titre calculation

HEK293T cells (2.5×10^6^) were seeded on 100-mm cell culture dish (Corning, USA #430167) 24 hours before transfection. Transfer lentiviral plasmids (2500 ng) were diluted in 200 µl serum-free medium together with 225 ng pCMV-VSV-G (Addgene #8454) and 2250 ng psPAX2 (Addgene #12260) plasmids. FuGENE 6 transfection reagent (Promega, USA #E2691) diluted in serum-free DMEM was added to the plasmid mixture at 1.5:1 (reagent: DNA) ratio. The mixture was incubated at room temperature for 30 minutes and added to cells dropwise. Cell media was refreshed after 16 hours of incubation and viral supernatant was collected 48 and 72 hours after transfection. Viral supernatant was filtered with 0.45 µm syringe filters (Corning, USA #431225) and precipitated with 50% poly(ethylene glycol) (Sigma-Aldrich, USA #25322-68-3) at a 4:1 (supernatant: PEG) ratio. After 24 hours of incubation at 4°C, viral particles were precipitated by centrifuge and the pellet was resuspended with 1 ml DPBS (Biowest, Franse #L0615) and stored at −80 °C until use. Viruses were titered and then used to transduce cells supplemented with 8 µg/ml protamine sulfate (Sigma-Aldrich, USA #P3369).

### Proliferation assay

Cell proliferation was measured by Cell-Titer Glo (CTG) Luminescent Cell Viability Assay (Promega, USA #G7570), MTS Colorimetric Cell Proliferation Assay (Promega, G3580), and manual cell counting with a hemocytometer. For CTG and MTS assays, cells were seeded in black 96-well plates (Corning, USA #3603) and transparent 96-well plates (Sarstedt, Germany #83.3924) respectively with 2000 cells/well. Cells were incubated for 96 hours and the proliferation rates were measured. For the CTG assay; cell medium was aspirated and a mixture of 40 µL medium + 4 µL Cell-Titer Glo reagent was added to each well and the relative luminescence was measured using BioTek Synergy H1 Hybrid Multi-Mode Reader (Agilent, USA). For the MTS assay; 20 µL MTS reagent was added to each well and the plates were incubated at 37 °C for 2 hours. Absorbance was recorded at 490 nm using BioTek Synergy H1 Hybrid Multi-Mode Reader. For manual cell counting, 100,000 cells/well were seeded in 6-well plates and cell numbers were counted with a hemocytometer after 96-hour incubation.

### Luciferase assay

Validated AR enhancers (750–850 bp) were selected for testing based on the HOXB13 ChIPseq and stratified as either “HOXB13 binding low” or “HOXB13 binding high” (Supplementary Table 2). They were assayed with a luciferase reporter assay as previously described (25). Briefly 293T cells were co-transfected with 100 ng reporter plasmid, 5 ng of Renilla, 100 ng pEZY_AR plasmid and either 100 ng pEZY_eGFP or pEZY_Flag_HOXB13 using TransIT-2020 (Mirus) and then plated in DMEM (Gibco) supplemented with 10% FBS (Fisher Scientific). 48 hours post-transfection, cells were treated with 10 nM of DHT or ETOH for 24 h. Firefly and Renilla luciferase activity were assayed by Dual-Glo Luciferase assay system (Promega). Relative activity was calculated by fixing the AR and eGFP transfected cells at 100%. All the experiments had a minimum of 4 biological replicas with 2 technical replicates in each experiment.

### *In vivo* animal studies

All in vivo experiments were approved by Koç University Animal Experiments Ethics Committee. LNCaP or PC3 cells stably expressing pLENTI CMV V5-Luc Blast were transduced with lentiCRISPRv2-HOXB13_sgRNA #1 or non-targeting sgRNA. LNCaP (3.5 x 10^6^ cells/animal) were mixed with Matrigel hESC-qualified Matrix (Corning, 354277) at a 1:1 (vol/vol) and injected subcutaneously into eight male nude mice. PC3 (1.5 x 10^6^ cells/animal) were directly injected subcutaneously in the flank side of the animal into male nude mice (n=15). For tetracycline-inducible experiments, PC3-luciferase cells were transduced with either Tet-pLKO_shHOXB13 #1 or Tet-pLKO_shFF. Following antibiotic selection, cells (1.5 x 10^6^ cells/animal) were injected subcutaneously into male nude mice (n=20). Once the tumors became palpable, 500 µg doxycycline was injected intraperitoneally into the animals on alternate days. Tumor volume was monitored weekly by caliper measurements. Upon completion of the study, tumors were excised and imaged.

### Chromatin immunoprecipitation-sequencing (ChIPseq)

ChIPseq was performed as previously described (26). Briefly, 5 x 10^6^ LNCaP and PC3 cells were cultured in RPMI supplemented with 10% FBS for 48 hours. Cells were crosslinked with 1% methanol-free formalin for 10 min at room temperature with gentle shaking. After lysis and nuclear enrichment, immunoprecipitation was carried out using HOXB13 (Cat # ab227879, Abcam), and SMARCD2 (Cat # ab221168, Abcam) -conjugated protein G Dynabeads (Cat # 10003D, Invitrogen) overnight at 4°C. After washing, the DNA was eluted and de-crosslinked at 65°C for 8 hours. The DNA was purified using Monarch DNA purification kit (Cat # T1030, NEB) and quantified by Qubit high-sensitivity DNA kit (Cat # Q33230, Invitrogen). DNA (5ng) was used for sequencing library construction using NEBNext Ultra II DNA Library Prep Kit for Illumina (Cat # E7645, NEB) according to the manufacturer’s instructions. The library was amplified for 10 cycles and quantified by Bioanalyzer (Agilent) and sequenced using Illumina NovaSeq 6000 with a minimum of 29M paired-end reads for each sample.

### Assay for Transposase-Accessible Chromatin using sequencing (ATACseq)

ATACseq was performed as described (27) with minor modifications. Briefly, 50,000 live DU145 cells expressing GFP or HOXB13 were lysed in 50 µL of cold lysis buffer (10mM Tris-HCl, pH 7.5, 10mM NaCl, 3mM MgCl2, 0.1% NP-40, 0.1% Digitonin, and 0.1% Tween-20) on ice for exactly 3 mins. The intact nucleus was tagmented with 2.0 µL of Tn5 Transposase (Cat # FC-121-1030, Illumina) and incubated at 37°C on a thermomixer at 1000 RPM. The DNA was purified by the Minielute reaction cleanup kit (Cat # 28206, Qiagen). The DNA was amplified with unique barcoded primers (Supplementary Table 1) and subjected to sequencing using Illumina Novaseq 6000 with a minimum of 30M reads for each sample.

### ChIPseq and ATACseq data analysis

Quality checks of raw and concatenated FASTQ files were done by FastQC (v0.11.9), and compared using MultiQC (v1.11) (28). Paired-end reads were aligned to the hg38 genome build using BWA (v0.7.17-r1188) (29). Duplicate alignments and alignments with a MAPQ score lower than 30 were removed. *ChIPseq:* Binding regions were called with MACS3 (v3.0.0a6) (30). Signal bedgraph files were generated using bedtools (v2.30.0) (31) and converted into bigwig files using bedGraphToBigWig (v4) tool. The signal was TMM normalized (32) with calculated normalization factors using edgeR (v3.36.0)(33). *ATACseq:* Mapped mitochondrial DNA was filtered (34) and accessible peaks were called using MACS3 (v3.0.0a6). *Patient ChIPseq:* Raw single end sequencing reads for patient ChIPseq samples were obtained from GSE130408 and aligned to the hg38 genome build using BWA (v0.7.17-r1188). Duplicate alignments, alignments with a MAPQ score lower than 30 and unmapped reads were removed. Binding regions were called with MACS2 (v2.1.1). All publicly available data is described in Supplementary Table 3.

### Motif analysis

FIMO (v5.3.0,(35)) was used to identify matches for HOCOMOCO v11 CORE motifs (36). Natural logarithm of q-values was calculated for each motif instance, and grouped by each factor for each set to average significance. Differences between average log q-values of each factor between set pairs were calculated and ranked.

### Single-cell RNAseq analysis

Analysis of published primary PCa single-cell RNAseq (GSE176031) was conducted as previously described (37).

### RNA-seq analysis

*AR* and *HOXB13* expression was quantified from publicly available RNAseq of CRPC (n=99) patient cohorts (38). Gene Set Variation Analysis (GSVA) was used to calculate the published AR (ARG10; *ALDH1A3, FKBP5, KLK2, KLK3, NKX3-1, PART1, PLPP1, PMEPA1, STEAP4, TMPRSS2*) signature and t-NEPC (NEG10; *C1QBP, CHGA, CHGB, CHRNB2, ELAVL4, ENO2, PCSK1, SCG3, SCN3A, SYP*) signature (40).

### Patient randomization

Clinical ATACseq (CRPC and NEPC: GSE156291, Primary PCa: (41)) and HOXB13 ChIPseq (GSE130408) were bootstrapped (1000x) to generate randomized genomic regions with the same size distribution as experimental ATACseq peaks and then intersected with clinical ChIP or ATAC narrowPeaks.

### Rapid Immunoprecipitation Mass spectrometry of Endogenous protein (RIME)

RIME was performed as previously described (42). Briefly, 2×10^7^ LNCaP, PC3, or 22Rv1 cells were crosslinked with 1% methanol-free mass spectrometry grade formalin for exactly 8 min at room temperature. After cell lysis and nuclear enrichment, immunoprecipitation was carried out using HOXB13 (Cat # ab2227879, Abcam) or IgG (Cat # AB-105-C, R&D Systems) -conjugated protein G Dynabeads (Cat # 10003D, Invitrogen) overnight at 4°C with rotation. The beads were rigorously washed for 10X with cold RIPA buffer (50 mM HEPES (pH 7.6), 1 mM EDTA, 0.7% (wt/vol) sodium deoxycholate, 1% (vol/vol) NP-40 and 0.5M LiCl) followed by 2X wash with cold 100mM ammonium bicarbonate in a protein low-bind tube. The beads were thoroughly dried and stored in −80°C till all the biological repeats were completed and ready for trypsin digestion.

### Label-free LC-MS/MS proteomics

On bead trypsin digestion was carried out in 50µL HEPES pH 8.0 with 1µg trypsin at 37°C overnight. The digested peptide concentration was determined using a Nanodrop/BCA assay, desalted with C18 TopTips (200µL), lyophilized by Centrivap, and dissolved at 0.7µg/µL with 0.1% formic acid. Analysis of peptides was carried out on an Orbitrap Fusion Lumos MS platform (Thermo Scientific) coupled to an Easy-nLC 1200 system (Thermo) using an in-house 100 µm ID x 100 cm column (Dr.Maisch 1.9µ C18) and EasySpray source (Thermo). The analytical column was equilibrated at 700 bar for a total volume of 8 μL and the injection volume was 2 μL. A gradient of mobile phase A (water and 0.1% formic acid) and B (80% acetonitrile with 0.1% formic acid) at 0.25µL/min, 2-27%B from 2-90 min followed by 27-40% B over 12 min, 40-95% B over 8 min and 10 min at 95%. Data acquisition was carried out using a data-dependent method with MS2 in the Orbitrap using a positive ion spray voltage of 2100, transfer tube temperature of 325°C, and default charge state 2. Survey scans (MS1) were acquired at a resolution of 120K across a mass range of 350 – 1500 m/z, with RF 30, an AGC target of 4e5, and a maximum injection time of 50ms in profile mode. For MS2 scans there was an intensity threshold of 5e4, charge state filtering of 2–5, dynamic exclusion 30 sec with 10ppm tolerances with a 1.2 m/z window before stepped HCD fragmentation of 31,33,37% using 7.5K resolution, auto mass range, AGC target 5e4 and maximum injection time of 124ms in centroid mode. All data files were processed with Protein Discoverer 2.5. Spectrum files were recalibrated and features were extracted with Minora. Searches were carried out with Sequest HT with SwissProt TaxID=9606 (v2017-10-25) with precursor mass tolerance 10ppm and fragment mass tolerance 0.01Da, carbamidomethyl static modification and K,M,P oxidation and S,T,Y Phos dynamic modification. Decoy database strict and relaxed FDR targets were 0.01 and 0.05 based on q value. Precursor quantification was intensity based on unique and razor peptides used, normalized by total peptide amount with scaling on all averages. The label-free quantification (LFQ) intensity from all biological replicates of HOXB13 and matched IgG controls were log2 normalized. The differential enrichment of protein in HOXB13-RIME over IgG-RIME was performed by NormalyzerDE using default parameters (43). Proteins with HOXB13/IgG with log2FC >2 and p<0.05 were considered significant. The functional role of significantly enriched proteins was determined by Metascape (44). Proteins identified from HOXB13 RIME study are listed in Supplementary Table 4.

### Proximity Ligation Assay (PLA)

To assess HOXB13-SMARCD2 interaction in prostate cancer cells PLA was carried out using Duolink® kit (Cat # DUO92101, Sigma Aldrich) according to the manufacturer’s instructions. Briefly, cells were fixed with 4% Formalin for 10 minutes at room temperature followed by permeabilization with 0.3% TritonX-100. The cells were incubated overnight at 4°C with rabbit anti-HOXB13 (Cat # 90944, Cell Signaling) or anti-IgG (Cat # AB-105-C, R&D Systems) antibody with mouse anti-SMARCD2 (Cat # sc-101162, SCBT). The protein-protein interactions were amplified with red fluorescence, and the signal was quantified by confocal microscopy (Olympus) under 60X magnification. HOXB13-SMARCD2 interactions in clinical samples were analyzed in a tissue microarray containing 122 prostate specimens (24 benign, 42 localized PCa, and 46 metastatic PCa) and 24 non-prostatic tissue specimens. To assess their interactions, Duolink® (Cat # DUO92012, Sigma Aldrich) was adapted from the manufacturer’s protocol to use on Ventana DISCOVERY Ultra autostainer as described (45). Briefly, antigen was retrieved at 91°C for 64 minutes. The tissues were incubated with anti-HOXB13 (Cat # 90944, Cell Signaling) and anti-SMARCD2 (Cat # sc-101162, SCBT) for 12 hours at room temperature. After ligation for 1 hour and amplification for 2 hours, the cells were stained with the nuclear staining solution provided in the kit. The protein-protein signal was scanned and scored digitally by a pathologist with Aperio image scope software (Leica Microsystems).

### Protein co-immunoprecipitation

pEzy-Flag-HOXB13 or pEzy-Flag-GFP was co-transfected with pEzy-myc-his-SMARCD2 in HEK293 cells. After 48 hours, cells were lysed with ice-cold IP lysis buffer (10mM Tris-HCl pH 7.4, 10mM NaCl, 1% Triton X 100, 1% NP-40). The lysate was incubated with anti-FLAG antibody (Cat # F1804, Sigma) conjugated magnetic beads (Cat # 1003D, Invitrogen) overnight at 4°C. After washing with IP buffer (10mM TRIS HCl pH 7.4, 1mM EDTA, 150mM NaCl, 1% TritonX 100) beads were boiled with 1X loading buffer. Protein was run on an SDS PAGE and probed with anti-his antibody (Cat # 2365, Cell Signaling Technologies).

### Immunohistochemistry (IHC)

A tissue microarray containing 139 specimens (35 untreated primary [hormone naive] tumors, 26 NHT tumors, 37 CRPCs, and 41 NEPC) was obtained from the Vancouver Prostate Center Tissue Bank. Except for CRPC samples that were obtained via transurethral resection of the prostate, all specimens were collected through radical prostatectomy. IHC staining was performed with mouse anti-HOXB13 monoclonal antibody (Cat # 90944, Cell Signaling) using a Ventana autostainer (Discover XT from Ventana Medical System, Tucson, AZ) with an enzyme-labeled biotin-streptavidin system and a solvent-resistant DAB Map kit (Ventana). HOXB13 staining intensity was scored by a clinically trained pathologist.

## RESULTS

### HOXB13 expression is independent of AR

While controversial, several studies have proposed that *HOXB13* gene expression is regulated by AR in PCa (17–19). To investigate this relationship, we quantified the expression of HOXB13 and AR in clinical PCa cohorts. We found that HOXB13 is highly expressed in primary PCa, CRPC, and AR-positive treatment-induced neuroendocrine prostate (t-NEPC) (**Figure 1A**). Similar to a recent study we observed that there was a loss or reduction of HOXB13 expression in most AR-negative t-NEPC clinical samples (4/7) (46). However, this was not a binary relationship and many AR-negative tumours (3/7) continued to express medium to high levels of HOXB13 (**Figure 1B**). With independent CRPC patients (38), HOXB13 mRNA expression was observed to correlate but not be found exclusively in AR+ tumours (**Figure 1C**). This was not limited to cell types as when we quantified *AR* and *HOXB13* in published single-cell RNA sequencing (scRNA-seq) of primary PCa (37), we found that while both *AR* and *HOXB13* were commonly co-expressed in luminal epithelial cells while both fibroblasts and endothelial cells solely express *AR* but not *HOXB13* (**Supplementary Figure 1**). When looking at the expression in different AR-low/negative CRPC tumour types, we found high expression of *HOXB13* in both neuroendocrine and stem-cell-like CRPC, but almost none in WNT-driven CRPC (**Figure 1D**). These results suggest that HOXB13 expression is not driven by AR activity. To explore this further, we then characterized HOXB13 expression following inhibition of AR in a primary PCa neoadjuvant cohort (**Figure 1E**). We found inhibiting AR with androgen deprivation therapy did not significantly decrease HOXB13 expression. Overall, these findings suggest that HOXB13 expression is not directly regulated by AR in PCa.

**Figure 1:**
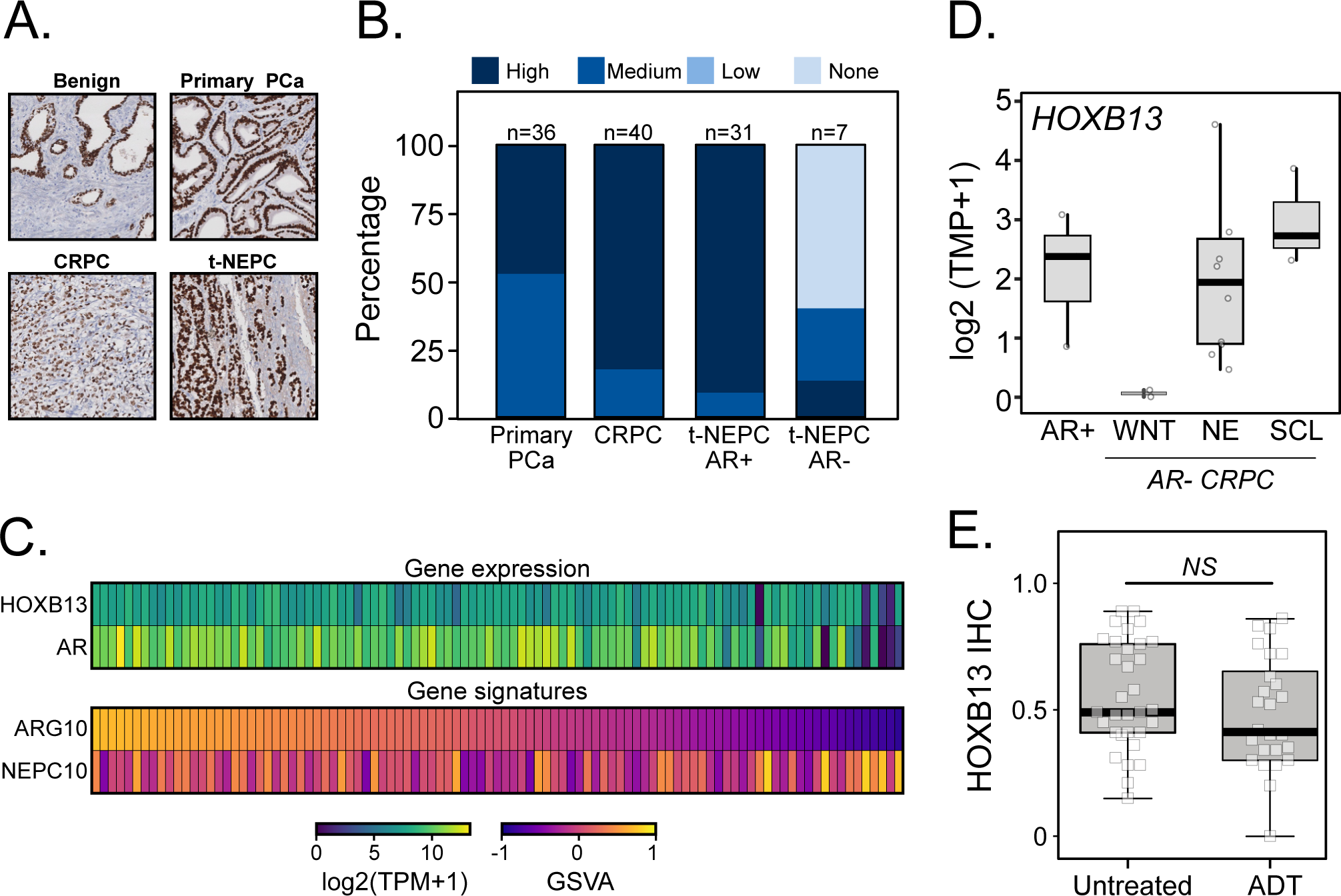
HOXB13 expression is independent of AR. **(A)** HOXB13 expression was evaluated by IHC in a tissue microarray containing primary PCa, CRPC, AR-positive (AR+) t-NEPC, and AR-negative (AR-) t-NEPC (n=114). **(B)** HOXB13 expression from the TMA IHC was graded by a clinical pathologist as high, medium, low, or no HOXB13 expression. HOXB13 expression was reduced, but still commonly found in AR-t-NEPC tumours **(C)** HOXB13 and AR expression was quantified from publicly available RNA-seq of laser-capture microdissected CRPC tumor tissue (n=99) and compared to published AR and NEPG gene signatures (ARG10 and NEPC10 respectively). (**D)** HOXB13 expression was compared in AR+ and AR-CRPC subtypes (50). High expression was observed in AR+, neuroendocrine and stem-cell like (SCL) CRPC while no expression was observed in WNT-driven CRPC. **(E)** HOXB13 IHC scored from PCa samples collected in a high-risk primary PCa neo-adjuvant clinical trial comparing untreated vs. androgen deprivation therapy (ADT). No change in HOXB13 was observed in untreated and ADT PCa samples.

### HOXB13 is selectively essential to PCa

Given the need for novel therapeutic targets that are independent of AR we then explored the potential impact of HOXB13 on PCa growth. We first knocked out HOXB13 *in vitro* with CRISPR/Cas9 using three separate sgRNAs in androgen-dependent PCa cell lines (LNCaP, VCaP, 22Rv1); androgen-independent AR-positive PCa cell lines (LNAI, LNCaP ABL); AR-negative PCa cell line (PC3); and non-PCa cell lines (293T, Hep3B). In all HOXB13 expressing PCa cell lines (**Supplementary Figure 1B**), CRISPR/Cas9 knockout significantly decreased cell proliferation with minimal effect on non-PCa cell lines (**Figure 2A; Supplementary Figure 2B**). Unexpectedly, this effect was independent of AR expression as the AR-negative PC3 cell line was the most sensitive to HOXB13 knockout. This phenotype was not due to guide RNA off-target effect as we observed similar effects on PCa proliferation with shRNA targeting HOXB13 (n=4) (**Figure 2B; Supplementary Figure 3A and B**). To expand on these PCa models, we quantified the *HOXB13* dependency from published genome-wide CRISPR screens (DepMap 22Q1;(47)). Similar to AR, we found that HOXB13 knockout was highly selective to PCa cell lines and did not significantly impact the proliferation of almost all non-prostatic cell lines (1059/1061) (**Supplementary Figure 4**). Expanding on these *in vitro* results we then tested the impact of HOXB13 on both *in vivo* xenograft initiation and growth. To characterize the impact of HOXB13 in tumour initiation, we first knocked out HOXB13 and generated subcutaneous xenografts in both AR-positive (LNCaP) and AR-negative (PC3) models. In these models, loss of HOXB13 caused a significant reduction of PCa engraftment with almost no tumour growth observed in any animals (**Figure 2C-D**). Next, we tested how HOXB13 impacted tumour proliferation in established tumours. Subcutaneous xenografts were generated with PC3 cells stably expressing inducible shRNA against either HOXB13 (shHOXB13) or firefly luciferase (non-targeting shCtrl) (**Supplementary Figure 5A and B**). Following the growth of a palpable tumour (volume ∼100 mm^3) shRNA expression was induced with exogenous doxycycline. No difference was observed in the growth of shHOXB13 and shCtrl xenografts before doxycycline treatment. However, following RNAi induction, those tumours expressing shHOXB13 showed a significant reduction in tumour size compared to the shCtrl group (**Figure 2E**). We next characterized the mechanism of cell death and found that loss of HOXB13 caused acute cell death by cellular apoptosis (**Figure 2F**). These results clearly demonstrate that HOXB13 is essential for both *in vivo* tumour initiation and growth.

**Figure 2:**
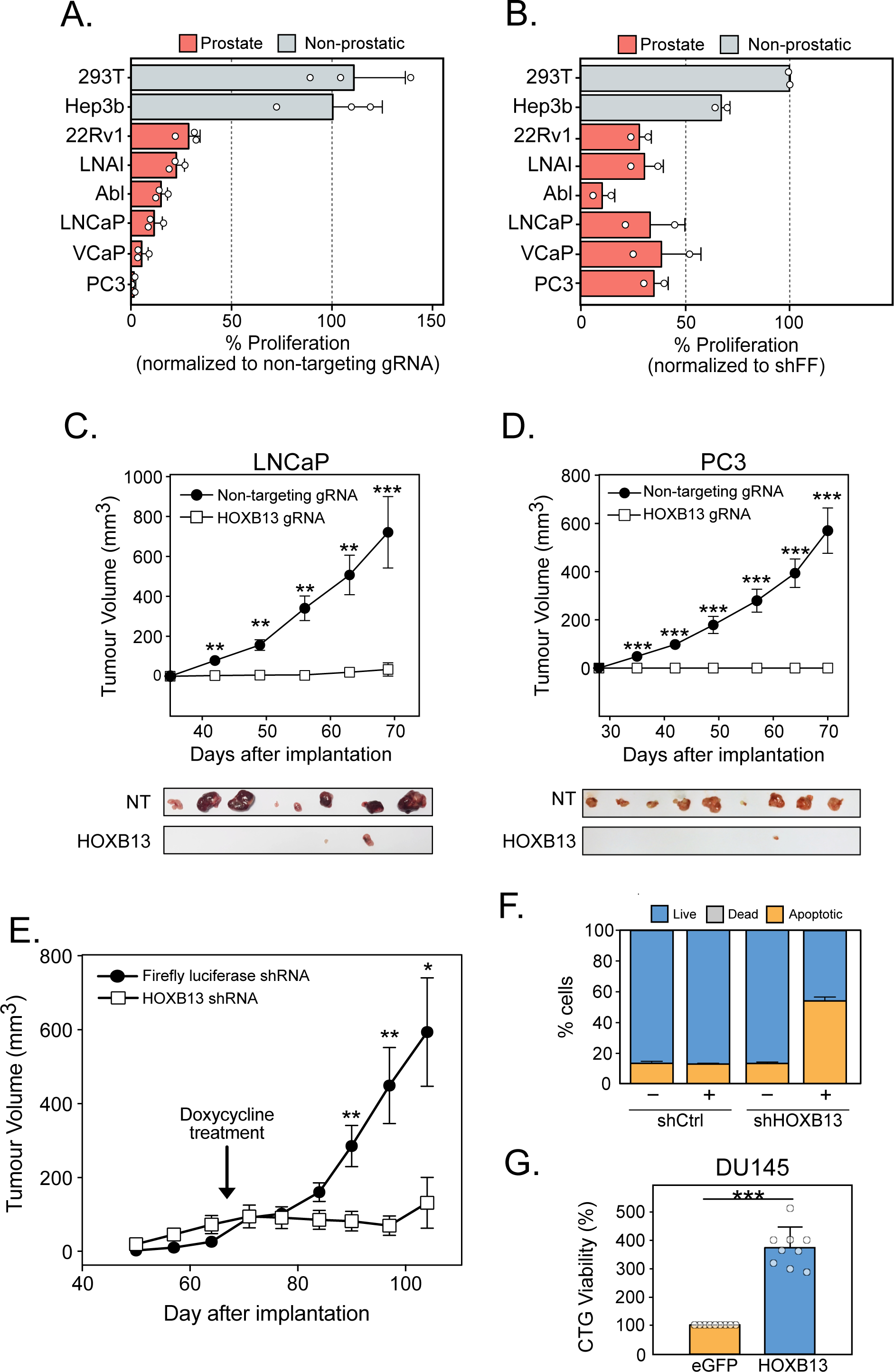
HOXB13 is selectively essential to PCa. **(A)** Proliferation assay in PCa and non-prostatic cell lines with HOXB13 knockout via CRISPR/Cas9 gene editing system (n=3; mean ± SEM); and **(B)** shRNA knockdown system showing the selective essentiality of HOXB13 for PCa (n=4 as biological replicates; mean ± SEM); **(C)** Tumor growth *in vivo* characterizing the effect of HOXB13 on PCa initiation in AR-dependent (n=8;mean ± SEM) and **(D)** AR-independent (n=15; mean ± SEM) subcutaneous xenografts; (**E)** Tumor growth *in vivo* characterizing the effect of HOXB13 in PCa progression (n=20; mean ± SEM unpaired two-tailed t-test). shCtrl and shFF were induced with doxycycline once the tumours became palpable; **(F)** Annexin V staining deciphering the effect of HOXB13 loss on apoptosis (n = 3; mean ± SEM); **(G)** Proliferation assay in HOXB13-null PCa (DU145) proving the essentiality of HOXB13 in PCa growth (n=9; mean ± SEM; unpaired two-tailed t-test). For all experiments an unpaired two-tailed t-test was used (ns *p>0.05, * p<0.05*, *\*\* p<0.01*, *\*\*\* p<0.001*)

Similar to previous work we found that *HOXB13* overexpression had a profoundly toxic effect on those cells dependent on HOXB13 (48) (**Supplementary Figure 6A**). Yet unexpectedly, we found that HOXB13 overexpression in DU145, an AR-negative PCa cell line that does not express HOXB13, significantly increased cellular growth (**Figure 2G**). This phenotype was confirmed in several orthogonal cell proliferation assays (**Supplementary Figure 6B-D**). This effect was specific to DU145 and overexpression of HOXB13 in a non-prostatic cell line (HEK293T) had no effect on cellular growth (**Supplementary Figure 6 E-G**). Together, our data show that HOXB13 is selectively essential in PCa initiation and growth independent of AR activity.

### HOXB13 has distinct binding in AR-positive and -negative prostate cancer

There is extensive work demonstrating that HOXB13 colocalizes at AR binding regions (ARBS) and affects AR-mediated transcription (14,15). However, as we demonstrated that HOXB13 is also essential for the proliferation in AR-negative cells, we reasoned that HOXB13 could potentially interact with alternative transcription factors in a cell-specific manner. To test this hypothesis, we characterized the HOXB13 cistrome in both AR-positive (LNCaP) and AR-negative (PC3) cells by chromatin immunoprecipitation followed by sequencing (ChIP-seq) (**Figure 3A**). We observed striking differences in the HOXB13 binding sites (HXBS) in these cell lines with minimal overlap in the HOXB13 cistrome (<10%) (**Figure 3A-C**). Strong HOXB13 binding was seen in LNCaP cells at *cis*-regulatory elements of AR-regulated genes, like *KLK3*, while no binding was seen in PC3 cells (**Figure 3B).** Ranked motif analysis showed that HXBS in AR-positive cells were enriched for AR and co-regulators motifs such as AR, FOXA1, and GATA2 while AR-negative PC3 cells were enriched for AP1 (JUN/FOS) transcription factor (**Figure 3D**). Comparison of the HOXB13 cistromes with published ChIP-seq studies using GIGGLE (49) showed that LNCaP HXBS significantly overlapped with AR cofactors including AR, ARD1A, FOXA1, and NKX31 (**Figure 3E**). In contrast, analysis of the HXBS in AR-negative PC3 cells showed significant overlap with binding sites of AP-1 (JUN/FOS) and ETV4 (50). The overlap with AP-1 binding sites was particularly interesting, as there has been recent work that defined a Stem-Cell Like (SCL) AR-low/negative CRPC subset that is driven by this transcription factor (50).

**Figure 3:**
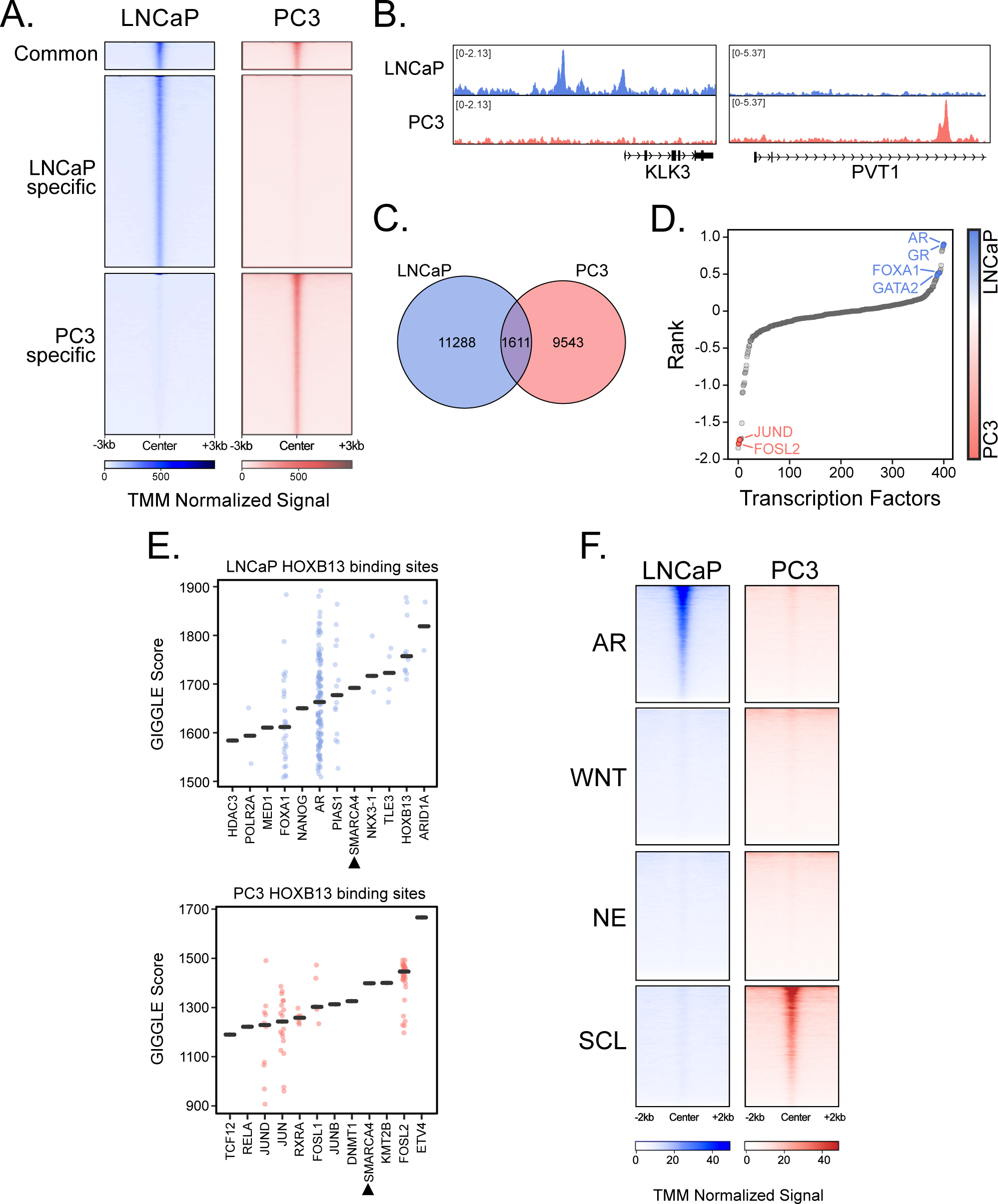
HOXB13 has a distinct cistrome in different stages of prostate cancer. **(A)** Heatmap of sorted TMM normalized HOXB13 ChIPseq signal in LNCaP and PC3 cells. **(B)** HOXB13 has distinct binding patterns in LNCaP and PC3 cells. **(C)** HOXB13 binding sites show minimal overlap (<10%) in LNCaP and PC3 cells. **(D)** Motifs ranked on their enrichment in LNCaP (Blue) and PC3 (Red) cells. **(E)** GIGGLE score of TFs shows SMARCA4 to be enriched at HOXB13 binding sites in both LNCaP (Left) and PC3 (Right) cells. **(F)** HOXB13 expression in CRPC organoid models with 4 different subtypes; CRPC-AR enriched in AR signature, CRPC-WNT enriched in WNT signature, CRPC-NE enriched in neuroendocrine (NE) signature, and high in NE markers, CRPC-SLC enriched in stem cell-like samples. **(G)** HOXB13 ChIPseq signal from LNCaP and PC3 cells on subtype-specific ATACseq peaks. LNCaP-HXBS is enriched in CRPC-AR and PC3-HXBS in CRPC-SLC with no overlap in their binding.

To explore the potential role of HOXB13 in AR-indifferent PCa, we mapped the PC3 HXBS cistromes to published ATACseq from these CRPC subtypes (**Figure 3F**). As expected, we saw a significant overlap with the LNCaP HOXB13 cistrome and the AR-dependent CPRC. Interestingly, the PC3 HXBS cistrome was highly enriched for the AP-1 driven AR-low/negative SCL CRPC suggesting that HOXB13 may interact with this transcription factor. Despite these differences in TF interactions, we reasoned that there potentially could be a common HOXB13 coregulator effector for all models. When we overlaid the motif and GIGGLE analysis from both AR-positive and -negative models we observed that both HOXB13 cistromes were commonly enriched for SMARC4A, a member of the Switch/Sucrose Non-Fermentable (SWI/SNF) chromatin remodeling complexes (**Figure 3E**). Overall, these results suggest that HOXB13 binding can potentially interact with different transcription factors in AR-positive and -negative PCa and there may be common effector proteins.

### Defining the HOXB13 interactome

HOX proteins regulate gene transcription in a tissue-specific manner through protein-protein interactions (51). Given the large differences we observed in DNA binding, we hypothesized that there could be a cell-specific HOXB13 interactome in AR-positive and -negative PCA. To characterize these differences, we performed HOXB13 Rapid Immunoprecipitation Mass Spectrometry (RIME) in both AR-positive (LNCaP and 22Rv1) and AR-negative PCa (PC3) models (n=3; **Figure 4A**). High HOXB13 peptide coverage was seen across all RIME biological repeats and cell lines (∼40%) with no HOXB13 peptides in IgG controls (**Figure 4B**). Using a conservative threshold (log2 fold change HOXB13/IgG>2 and *p*<0.05), we identified 73, 123, and 153 high-confidence HOXB13-interacting proteins in LNCaP, 22Rv1, and PC3 cells, respectively (**Figure 4C; Supplementary Table 4)**. While each cell line had a distinct HOXB13-interactome, there were marked differences between AR-positive and -negative cell lines. Specifically, most HOXB13-interacting proteins were commonly found in both LNCaP and 22Rv1 including many components of the AR-transcriptional complex such as AR, FOXA1, and NKX3-1 (**Figure 4C**). In contrast, the HOXB13 interactome in AR-negative PC3 was markedly different from selective AP-1 interactions (cJUN). Despite the unique HOXB13-transcription factor interactions in AR-positive and -negative models, we reasoned that there may be a common effector function that drives HOXB13 activity. Metascape analysis for biological functional enrichment of HOXB13 interacting proteins showed significant enrichment for RNA splicing and chromatin modifications in all the cells (44) (**Figure 4E**) including several members of the SWI/SNF chromatin remodeling complex (SMARCA4, SMARCA2, SMARCE1, ARID1A, SMARCD2, and PBRM1). This modular complex is a key regulator of nucleosome positioning and contains several core subunits, ATPase catalytic subunits, and variable subunits that when combined provide a distinct identity to each complex (52). In addition to various subunits identified in our RIME studies, SMARCD2 was found to interact with HOXB13 in all cells (**Figure 4C**). As previous studies have suggested HOXB13 can function as a pioneer factor and preferentially bind to methylated-CpG regions (53), we reasoned that HOXB13 activity can be mediated by this chromatin-modifying complex.

**Figure 4:**
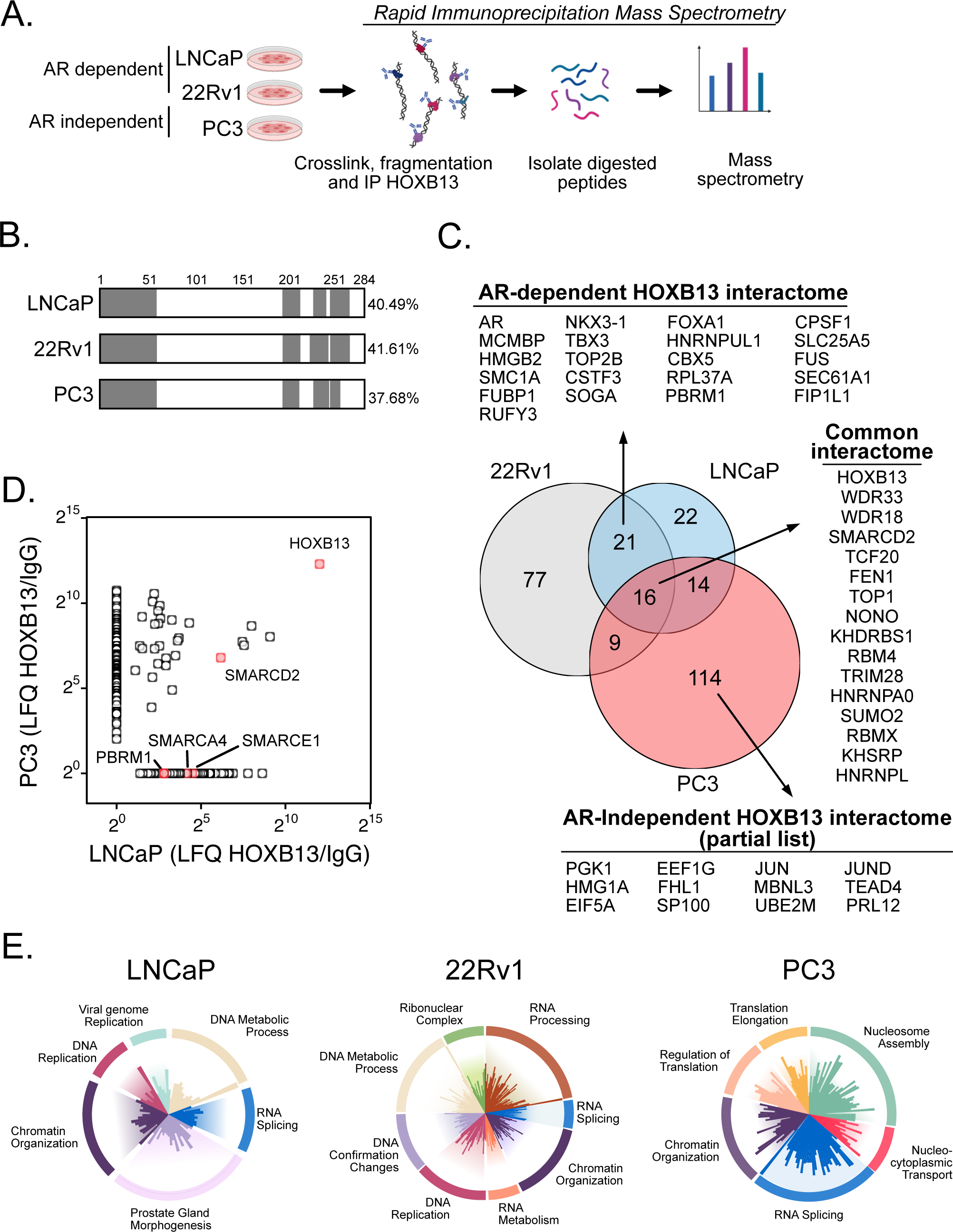
HOXB13 has a unique interactome in AR-dependent and AR-independent models. **(A)** Schematic representation of HOXB13-RIME. **(B)** Average peptide coverage of HOXB13 from individual RIME experiments in different cell lines. **(C)** HOXB13-interactome shows minimum overlap in AR-dependent and AR-independent PCa models. More than 50% of the H OXB13-interacting proteins are common to both LNCaP and 22Rv1 cells and are known AR-coregulators. **(D)** Many of the HOXB13-interacting proteins in AR-positive (LNCaP) and AR-negative (PC3) were markedly distinct. Members of SWI/SNF complex and HOXB13 are highlighted in red. Only **(E)** HOXB13-interacting proteins identified by RIME in LNCAP, 22Rv1, and PC3 cells. The proteins are grouped based on their biological functions and each bar represents the average Z-normalized LFQ intensities.

### Characterizing the HOXB13-SMARCD2 interaction

To validate these interactions between HOXB13 and SMARCD2, we performed *in-situ* proximity ligation assay (PLA) in multiple PCa cell lines (LNCaP, 22Rv1, PC3, and DU145). A strong PLA signal was seen in all the cells except DU145, which does not express endogenous HOXB13 (**Figure 5A and B**). When we overexpressed HOXB13 in DU145 cells we could observe an interaction with SMARCD2 (**Figure 5C**). The HOXB13-SMARCD2 interaction was confirmed by co-immunoprecipitation (CoIP) in cells exogenously expressing Flag-HOXB13 and Myc-His-SMARCD2 (**Figure 5D).** To determine if SMARCD2 co-occupies HOXB13 binding sites in prostate cancer cells we conducted SMARCD2 ChIP-seq (**Figure 5E).** We observed a significant overlap between HOXB13 and SMARCD2 cistromes in LNCaP cells. Interestingly, the HOXB13-SMARCD2 overlapping sites were significantly enriched at AR binding sites as compared to the HOXB13 alone sites (**Figure 5E and 5F**). Next, to test if HOXB13-SMARCD2 interactions were stabilized by AR binding we tested the PLA in LNCaP cells treated with androgens. We observed significantly higher PLA signals in those cells treated with DHT compared to EtOH (**Figure 5G; Supplementary Figure 7**). To characterize the potential impact of this on AR activity we tested the role of HOXB13 at validated AR enhancers (25). We found that HOXB13 significantly increased AR activity at HXB13 sites compared to unbound regions. (*p*<0.001; **Supplementary Figure 8**). Next, to understand the implications of HOXB13-SMARCD2 interactions for the clinical development and progression of prostate cancer, we analyzed the expression of HOXB13 and SMARCD2 in PCa cohorts. We found that HOXB13 and SMARCD2 expression correlated during the malignant transformation of the prostate to localized PCa (**Figure 6A**). To assess the HOXB13-SMARCD2 interactions in PCa, we performed PLA on clinical samples and found that HOXB13-SMARCD2 interactions occurred only in the prostate but not in non-prostate tissue (**Figure 6B and C**). This HOXB13-SMARCD2 interaction was observed in almost all clinical samples regardless of metastatic potential, tumour, node and metastasis (TNM) grading, or Gleason score (**Supplementary Figure 7C-E**). Taken together these results suggest that while HOXB13 interacts with unique transcription factors in AR-positive and -negative PCa, interactions with SMARCD2, a member of the SWI/SNF complex, offer a common effector function.

**Figure 5:**
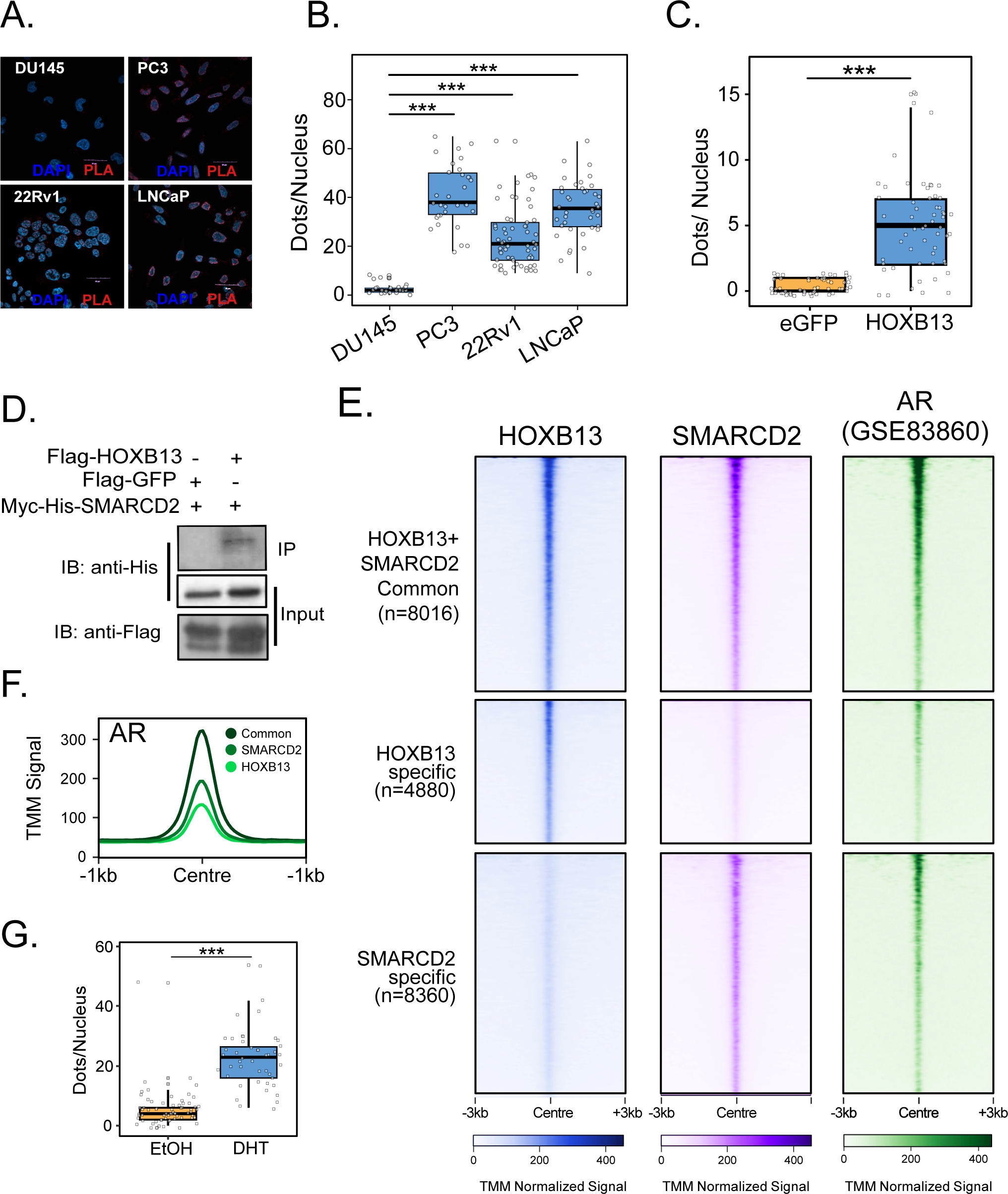
SMARCD2 interacts with HOXB13. **(A)** Representative confocal image of HOXB13-SMARCD2 interactions quantified by in-situ PLA. **(B)** HOXB13-SMARCD2 interactions were quantified manually and represented as dots/nuclei. The box plot represents results from 3 independent experiments (4 random 100X microscopy fields per experiment) and each dot represents an individual nucleus. **(C)** DU145 cells were transfected with either eGFP or HOXB13. In situ PLA shows significant HOXB13-SMARCD2 interactions only in DU145 cells expressing HOXB13. **(D)** CoIP in HEK293 cells co-transfected with Myc-His-SMARCD2 and Flag-HOXB13 or Flag-GFP. IP with anti-Flag antibody was probed with anti-His antibody. **(E)** LNCaP were cultured in androgen-deprived conditions for 72 hours and treated with 10nM of DTH or EtOH (Vehicle) for 4 hours. In situ PLA shows HOXB13-SMARCD2 interactions stabilized by androgen. **(F)** Heatmap of sorted TMM normalized ChIPseq peaks of HOXB13, SMARCD2, and AR in LNCaP cells. SMARCD2 co-occupies HXBS. For all data ns *p>0.05, * p<0.05*, *\*\* p<0.01*, *\*\*\* p<0.001*)

**Figure 6:**
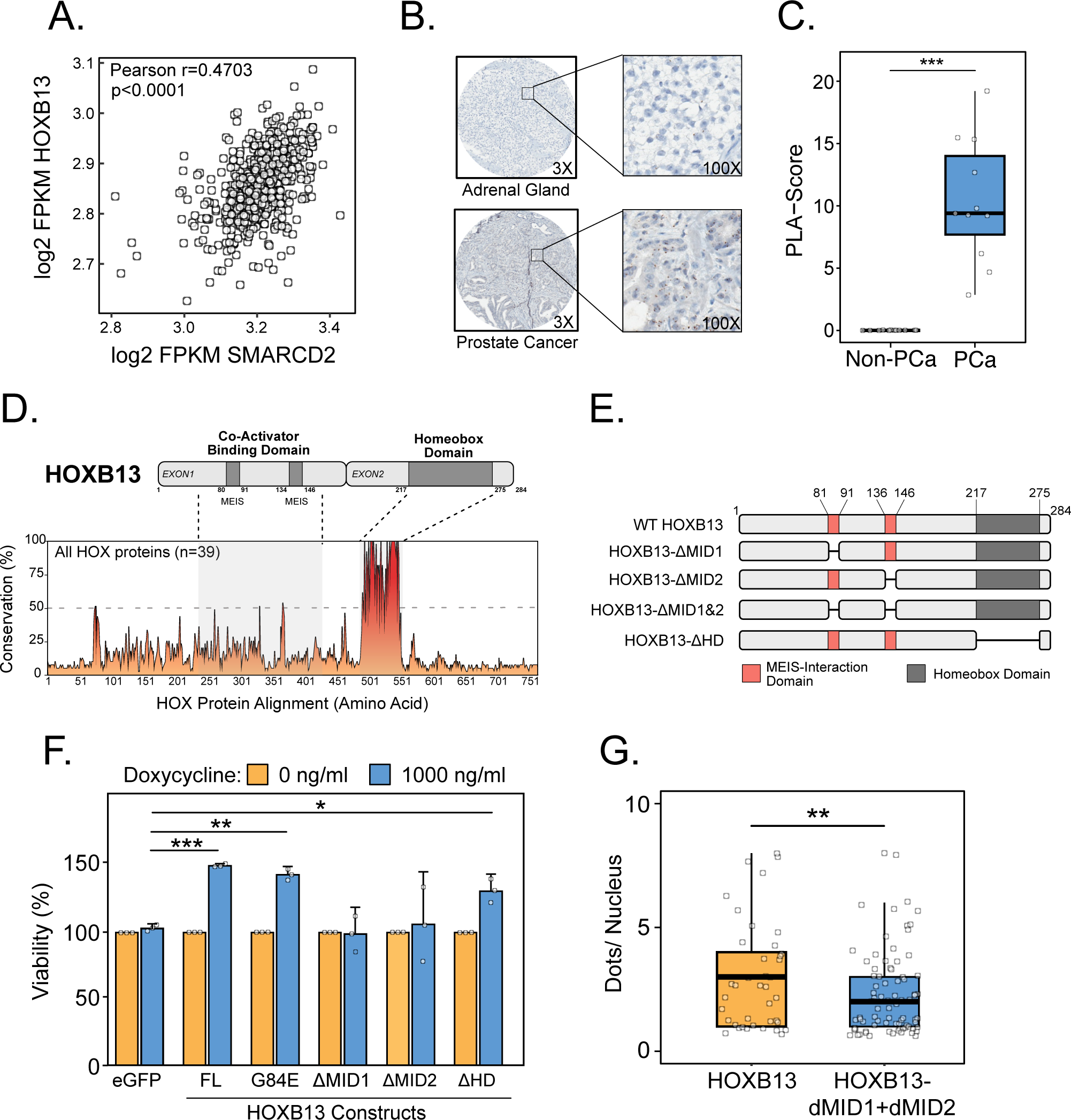
HOXB13 interacts with SMARCD2 through MEIS interaction domains and is clinically relevant. **(A)** Expression of HOXB13 and SMARCD2 is significantly increased in PCa compared to Benign tissue specimens in the TCGA cohort. **(B)** Expression of HOXB13 and SMARCD2 is significantly correlated in the TCGA-PCa cohort. **(C)** HOXB13-SMARCD2 was quantified by PLA in TMA consisting of normal and cancer samples from both prostate and non-prostate tissue. Representative images show interactions only in PCa (Bottom) but not in the adrenal gland (Top). (**D)** Quantification of PLA shows a significant increase in HOXB13-SMARCD2 interactions in PCa compared to non-PCa tissue specimens. **(E)** Schematics of various constructs with deleted HOXB13 functional domains. **(F)** Proliferation assay in DU145 with HOXB13 tetracycline-inducible overexpression revealing the essentiality of MEIS-interaction domains for HOXB13 to function (n=3; mean±SEM). **(G)** DU145 expressing FL-HOXB13 shows significantly more interactions with SMARCD2 compared to those expressing = both MID1 and MID2 deleted HOXB13 (HOXB13 -dMID1+dMID2) quantified by in-situ PLA. For all data ns *p>0.05, * p<0.05*, *\*\* p<0.01*, *\*\*\* p<0.001*.

### HOXB13 interacts with SMARCD2 through MEIS-interaction domains

Given the common HOXB13-SMARCD2 interaction, we hypothesized that this protein binding may be required for downstream activity. The HOXB13 gene encodes two distinct co-activator binding domains, MEIS-interaction domain-1 (MID1) and MEIS-interaction domain-2 (MID2), and a highly conserved DNA binding homeobox domain (HD) (**Figure 6D**). As HOXB13 overexpression increased proliferation of DU145 (**Figure 2G**), we conducted a gain-of-function experiment with distinct HOXB13 mutations/deletions including; G84E substitution (an SNP commonly seen in heritable PCa), ΔMID1, ΔMID2, and ΔHD (**Figure 6E**). No significant differences were observed in mutant HOXB13 protein expression (**Supplementary Figure 6E**). When we tested the effects of these mutants on the cell proliferation rates using tet-inducible expression systems, we found that HOXB13-ΔMID1 and HOXB13-ΔMID2 constructs did not increase DU145 cell proliferation while all other constructs (G84E and ΔHD) increased cell growth at a rate similar to WT HOXB13 (**Figure 6F**). To determine if the SMARCD2-HOXB13 interaction occurs through the MID1 and MID2 domains, we conducted HOXB13-SMARCD2 PLA in DU145 cells and observed a significant reduction in HOXB13-SMARCD2 interactions in those cells expressing HOXB13-ΔMID1+ΔMID2 (**Figure 6G**). Together, our results indicate that HOXB13 interacts with SMARCD2 through its MID1 and MID2 and these domains are essential for HOXB13-mediated activity.

### HOXB13-mediated recruitment of SMARCD2 increases chromatin accessibility

As SMARCD2 is a mSWI/SNF complex subunit, we reasoned that HOXB13 interactions could potentially alter the local chromatin accessibility. When we characterized chromatin accessibility with ATAC-seq in DU145 cells expressing either eGFP or HOXB13, we observed that HOXB13 expression caused a broad increase in chromatin accessibility at already formed euchromatin (**Figure 7A**). At the HOXB13-enriched sites (top 5%), we saw an increase of motifs associated with HOXB13, AR and FOXA1 (**Figure 7B**) and significant overlapped with clinical HOXB13 binding sites as compared to random sites (Mann-Whitney U *p*=1.5e-13) (**Figure 7C**). To evaluate the clinical relevance of HOXB13-enriched regions, we compared these results with clinical ATAC-seq from both primary PCa (41), CRPC and NEPC (54) (**Figure 7D**) and observed a significant overlap in HOXB13-enriched sites as compared to either eGFP-enriched or random sites in both primary PCa (*p*=7.3e-9) or CRPC (*p*=0.004). We did not see this same significance in AR-negative NEPC potentially due to the low number of patients and altered HOXB13 cistrome found in AR+ tumours. To determine if the HOXB13-mediated changes in chromatin accessibility occurred via SWI/SNF recruitment we characterized how SMARCD2 knockdown altered ATACseq in HOXB13 overexpressing cells (**Figure 7E**). Supporting the proposed mechanism, chromatin accessibility of the HOXB13-enriched regions was found to significantly decrease by SMARCD2 knockdown (**Figure 7F**). Together, these findings suggest that HOXB13 alters chromatin accessibility through SMARCD2-mediated activity.

**Figure 7:**
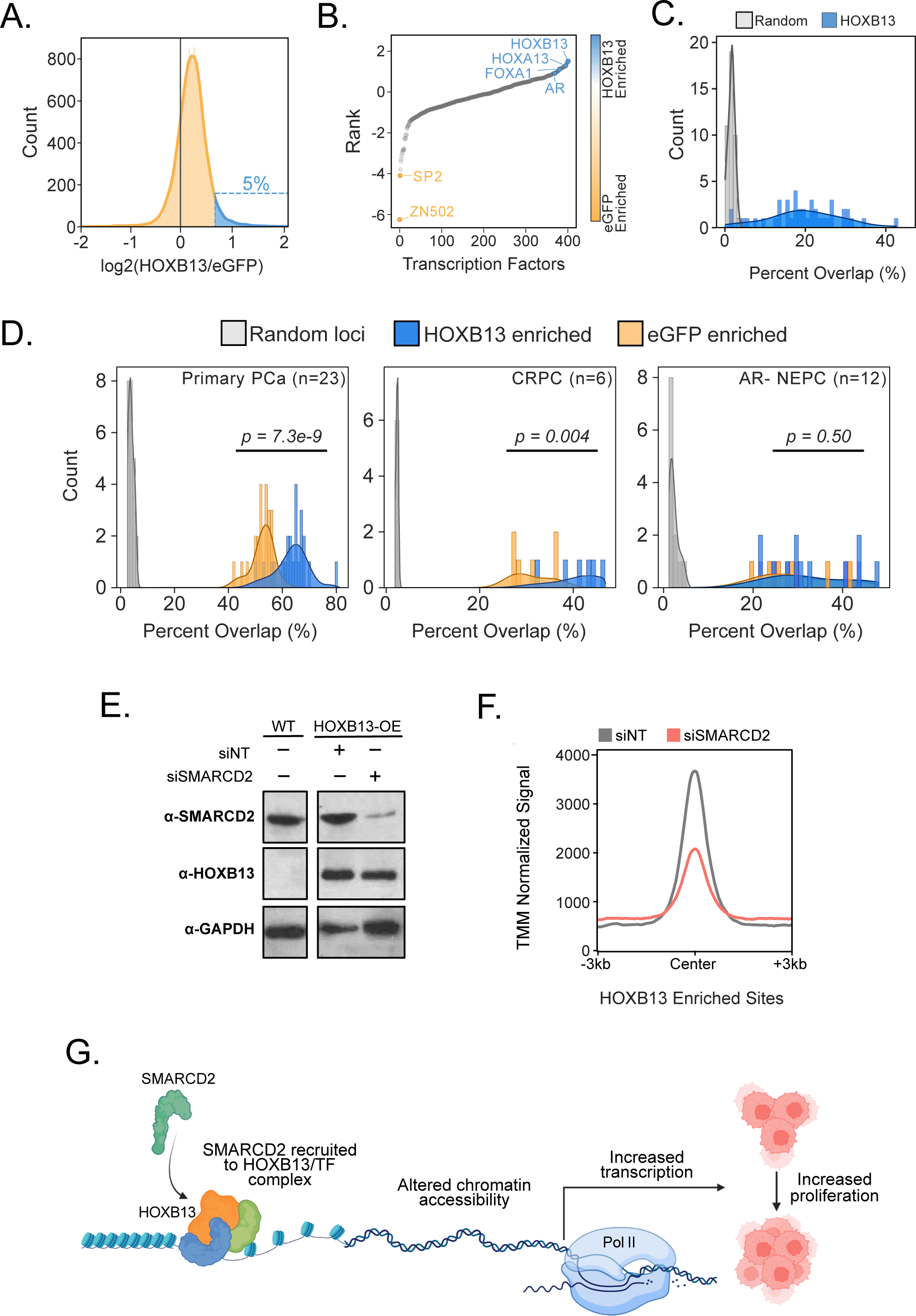
HOXB13-mediated recruitment of SMARCD2 increases chromatin accessibility. **(A)** ATACseq was performed in DU145 cells expressing with eGFP or FL-HOXB13. ∼ 5% of the accessible chromatin regions (highlighted in blue) are high-confidence HOXB13-specific open chromatin. **(B)** HOXB13-specific open chromatin is enriched for motifs from the HOX family and known AR-cofactors. **(C)** A significant overlap was observed for the HOXB13-enriched open chromatin regions with clinical HOXB13 binding sites (GSE130408) compared to random regions. **(E)** HOXB13-enriched sites are significantly enriched over the random or eGFP at the open-chromatin region in primary PCa (left panel), or CRCP (middle panel) but not in AR negative NEPC tissue specimens (right panel) **(E)** siRNA-mediated knock-down of SMARCD2 reduces the expression of SMARCD2 in HOXB13-expressing DU145 cells. No change in HOXB13 expression was observed. **(F)** Loss of SMARCD2 shows a marked reduction in ATAC-seq signal in HOXB13-expressing DU145 cells. (**G)** Schematic representation of the mechanism of HOXB13 in PCa, through the recruitment of SMARCD2 alters chromatin accessibility and increases the transcription of target genes to promote PCa growth.

## DISCUSSION

Despite the approval of several life-prolonging drugs in advanced PCa, resistance invariably emerges with most patients developing lethal CRPC. There is a clear clinical need for new therapeutics to control CRPC that is independent of AR. In this work, we demonstrate that HOXB13 is highly selective for tumor growth and proliferation of both AR-positive and -negative models. Strikingly, inhibiting this critical pathway has no significant effect on non-prostatic cell lines. Despite the common dependency, we show that HOXB13 PCa-specific cistromes and transcription factor interactions are markedly distinct between AR-positive and -negative PCa. Our work suggests that HOXB13 in SCL-CRPC interacts with AP-1 to drive cancer growth. Yet despite these differences in transcription factor interactions, the HOXB13 complex commonly interacts with SMARCD2 through the MEIS-interaction domains. Despite its role in increased PCa frequency, the common germline variant of HOXB13, G84E, was not associated with HOXB13 - SMARCD2 interaction suggesting that this is independent from our observed phenotype. Recruitment of SMARCD2 via HOXB13 alters chromatin accessibility and increases tumor progression in prostate cancer (**Figure 7G**). Importantly, this interaction is highly specific to prostate tissue and may offer a potential pharmacological target in CRPC.

Although several studies have characterized the proliferative effect of HOXB13 in prostate cancer, its role has been controversial (11,14,15,20–22). Our work shows that HOXB13 is selectively essential for both *in vitro* and *in vivo* PCa growth. While speculative the rapid kinetics of this effect (∼4 days) suggests that, in AR+ PCa cells, this was not solely due to inhibition of AR-mediated transcription as the cytotoxic effect of antiandrogens or AR-degraders occurs over a longer time frame (55,56). However, one challenge of this robust phenotype was that we had difficulty characterizing the impact of HOXB13 knockout on gene expression. We generated a HOXB13 knockout population but found that these resistant cells had a markedly altered morphology and growth compared to parental cells. Recent studies suggest that these AR+ HOXB13-null cell lines have distinct metabolic activity of action that occurs through loss of HDAC3 interactions (20). Notably, our findings represent the acute impact of HOXB13 inhibition in PCa models. Similar to published work we observed that HOXB13 expression can remain in AR-negative CRPC (46). However, in these patients, we demonstrate that HOXB13 remains active through AP-1 interactions. Given the importance of this transcriptional factor in double-negative CRPC (negative for both AR and NEPC markers) (50) this raises the intriguing possibility of selectively treating these patients by targeting HOXB13. However, further work is needed to elucidate the relationship between AP-1 and HOXB13 in double-negative CRPC.

## ACKNOWLEDGEMENTS

This work was supported by funding from the Canadian Institute of Health Science and TUBITAK (221Z116). SL was partially supported by a Prostate Cancer Foundation British Columbia PDF fellowship. The authors gratefully acknowledge the use of the services and facilities of the Koç University Research Center for Translational Medicine (KUTTAM).

