## Supplementary Figure for "HOXB13 alters chromatin accessibility in prostate cancer through interactions with the SWI/SNF complex"

### Supplementary Figure 1

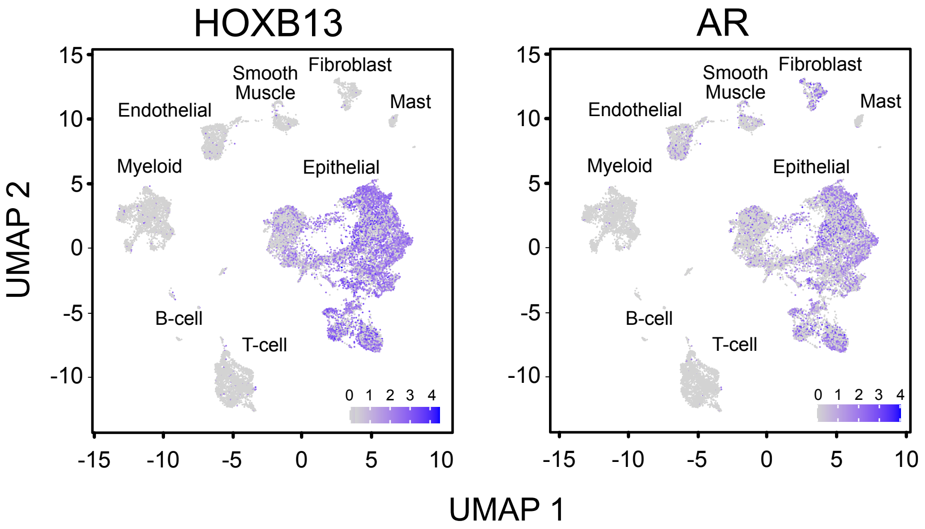


**Supplementary Figure 1:** Expression of AR and HOXB13 in single cell RNAseq from publically available primary PCa samples. UMAP projection of the major cell types identified by scRNA seq in a combined dataset of 21,743 cells from PCa specimens (GSE176031).

### Supplementary Figure 2

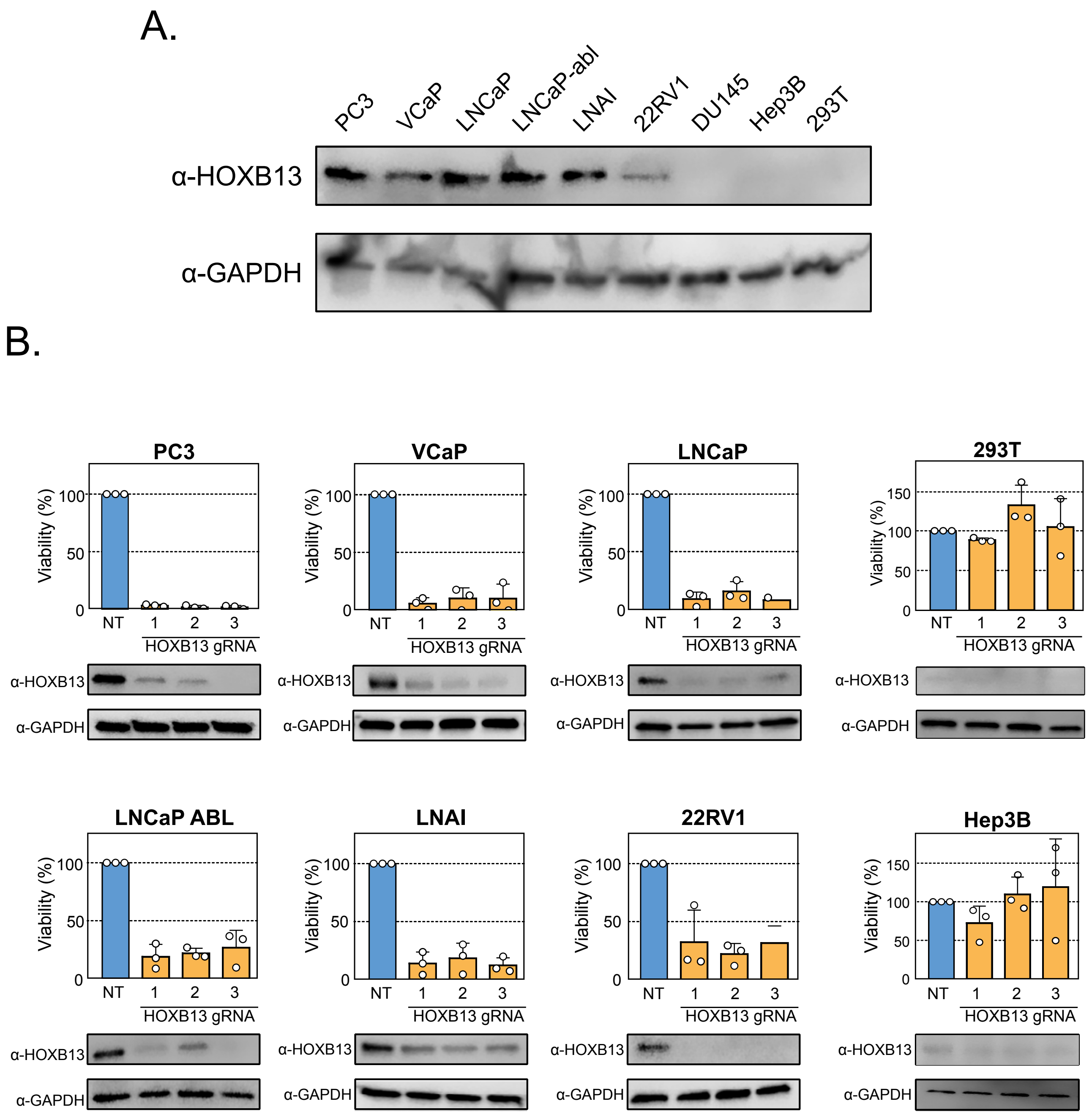


**Supplementary Figure 2: (A)** HOXB13 expression in different PCa and non-prostatic cell lines**. (B)**Proliferation assay for CRISPR/Cas9-mediated HOXB13 silencing in PCa (PC3, VCaP, LNCaP, LNCaP-ABL, LNAI, 22RV1) and non-prostatic cell lines (293T, Hep3B)(n = 3 biological replicates; Mean ± SEM). Western Blot validation of HOXB13 knockout with the individual gRNA is shown under the cell bar plot.

### Supplementary Figure 3

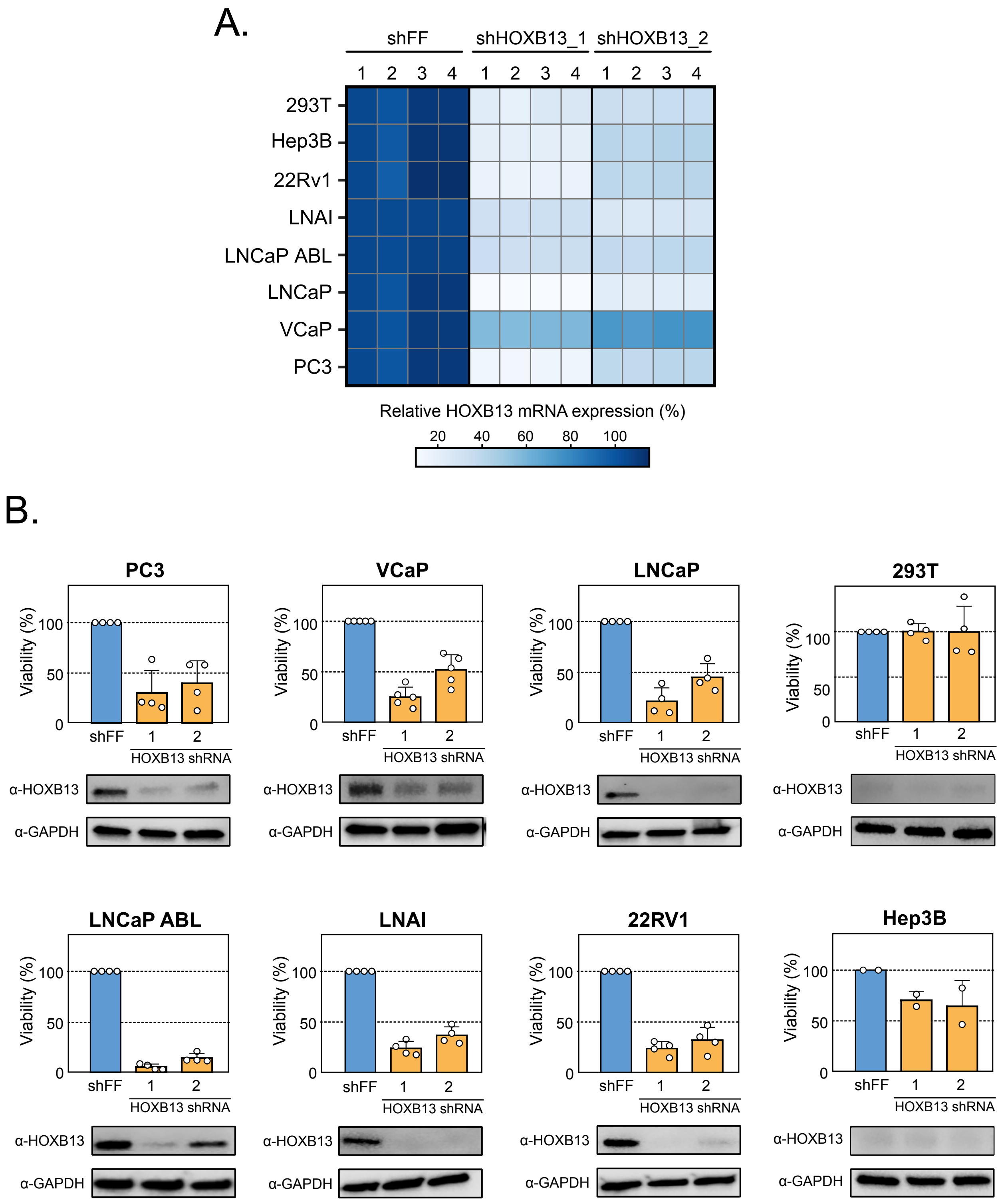


**Supplementary Figure 3:** Effect of HOXB13 shRNA on PCa (PC3, VCaP, LNCaP, LNCaP-ABL, LNAI, 22RV1) and non-prostatic cell lines (293T, Hep3B). **(A)** Quantification of HOXB13 expression by RT-PCR following shRNA treatment; **(B)** Proliferation assay for shRNA-based HOXB13 silencing (n = 4 biological replicates; Mean ± SEM). Western Blot validation of HOXB13 silencing with the individual shRNA is shown under the cell bar plot.

### Supplementary Figure 4

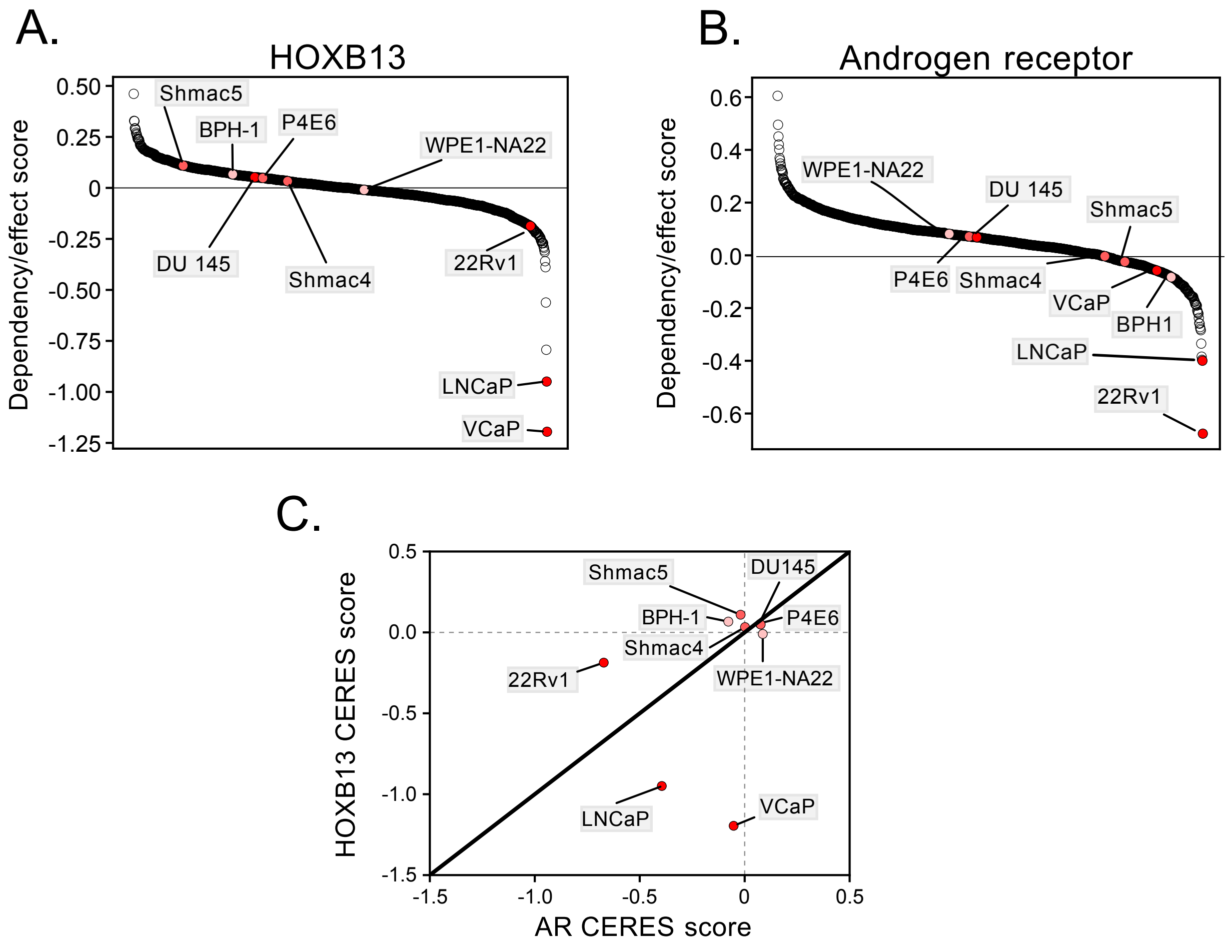


**Supplementary Figure 5:** *HOXB13* **(A)** and *AR* **(B)** CERES essentiality score across 1070 cells from Depmap 22Q1 release. Benign prostatic (light red) and prostate cancer (dark red) cell lines were selectively highlighted. **(C)** All prostate cell lines were compared for both *AR* and *HOXB13* essentiality.

### Supplementary Figure 5

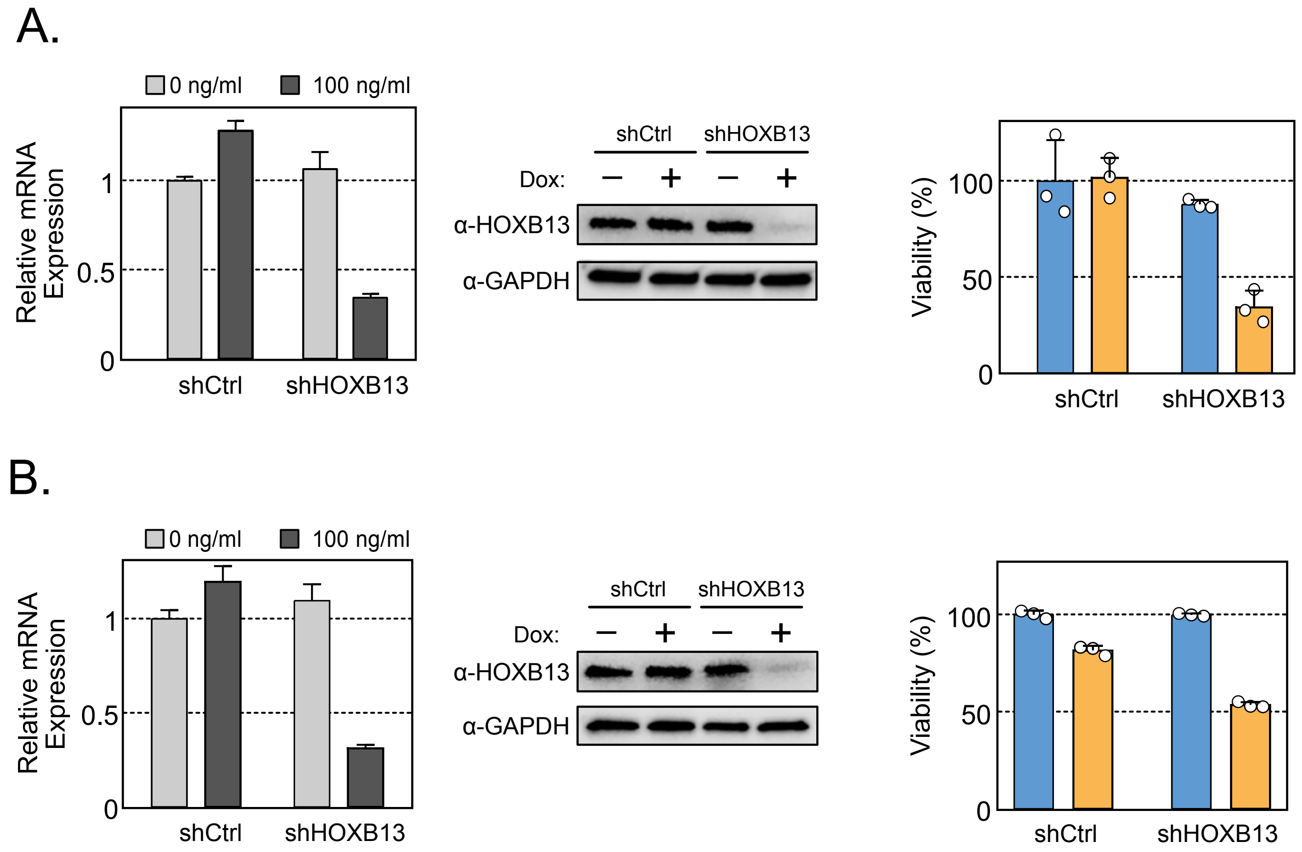


**Supplementary Figure 4:** RT-PCR and Western Blot validation, and proliferation assay of tetracycline-inducible shRNA targeting HOXB13 in **(A)** LNCaP; and **(B)** PC3 cell lines showing the silencing effect and decreased viability (n = 3 biological replicates; Mean ± SEM).

### Supplementary Figure 6

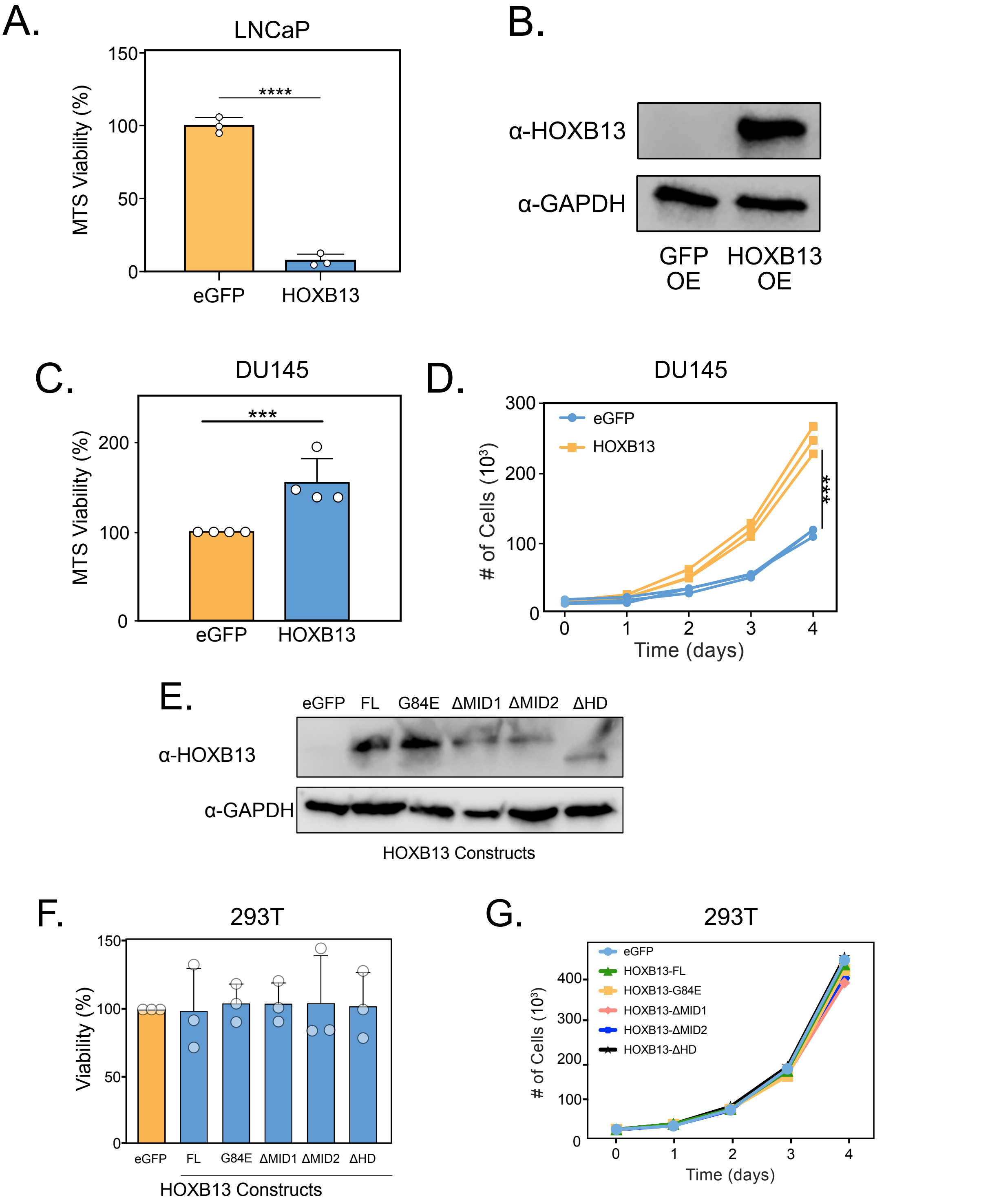


**Supplementary Figure 6:** Effect of HOXB13 expression on PCa and non-prostatic cell lines. **(A)** LNCaP cells were transfected with either CMV-eGFP or -HOXB13. Viability was quantified by MTS proliferation assay (n=3).**(B)** Western Blot of HOXB13 overexpression in HOXB13-null DU145 PCa cell line; **(C)** MTS proliferation assay (n=4); and **(D)** Manual cell counting quantifying proliferation rate in DU145 cells expressing either eGFP or HOXB13. No similar increase in proliferation was observed in non-prostatic 293T cells transfected with HOXB13 constructs and grown over 96 hrs. **(E)** Western Blot of HOXB13 overexpression in 293T cells (FL, G84E substitution, MID1 deletion, MID2 deletion, and HD deletion) in 293T cell lines. Proliferation was quantified by either manual cell counting **(E)** or MTS proliferation assay (n = 3)**(F)**. All data is shown as mean ± SEM; Unpaired two-tailed u-test was used to determine statistical significance

### Supplementary Figure 7

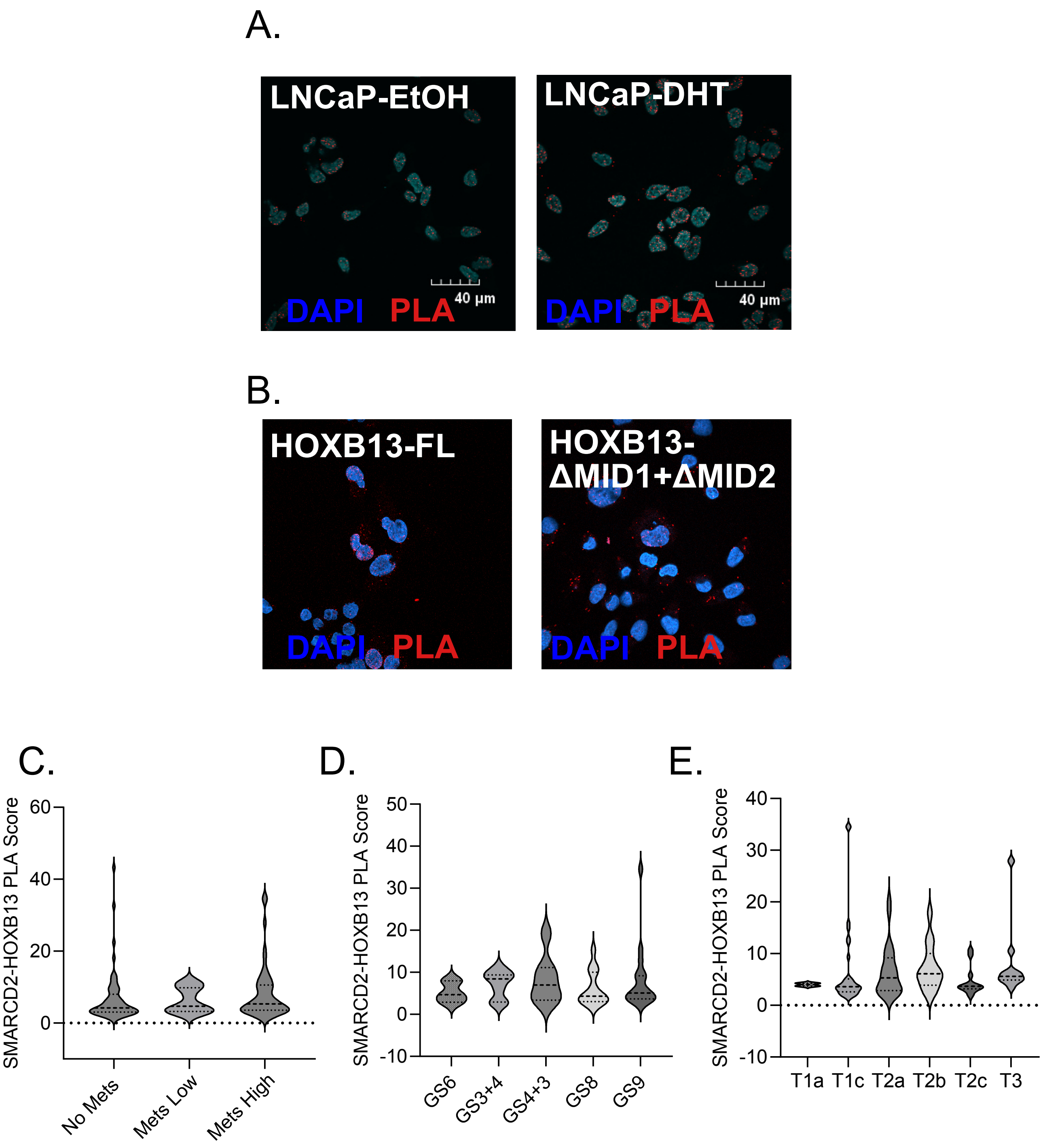


**Supplementary Figure 8:** Representative confocal image of HOXB13-SMARCD2 interactions quantified by in-situ PLA. **(A)** LNacP were cultured in androgen-deprived conditions for 72 hours and treated with 10nM of DTH or EtOH (Vehicle) for 4 hours. **(B)** DU145 were transfected with either FL-HOXB13 or MID1 and MID2 deleted HOXB13 (HOXB13-dMID1+dMID2). SMARCD2-HOXB13 PLA score from clinical PCa samples stratified by either (**C**) metastasis, **(D)** Gleason grade or **(E)** TNM staging.

### Supplementary Figure 8

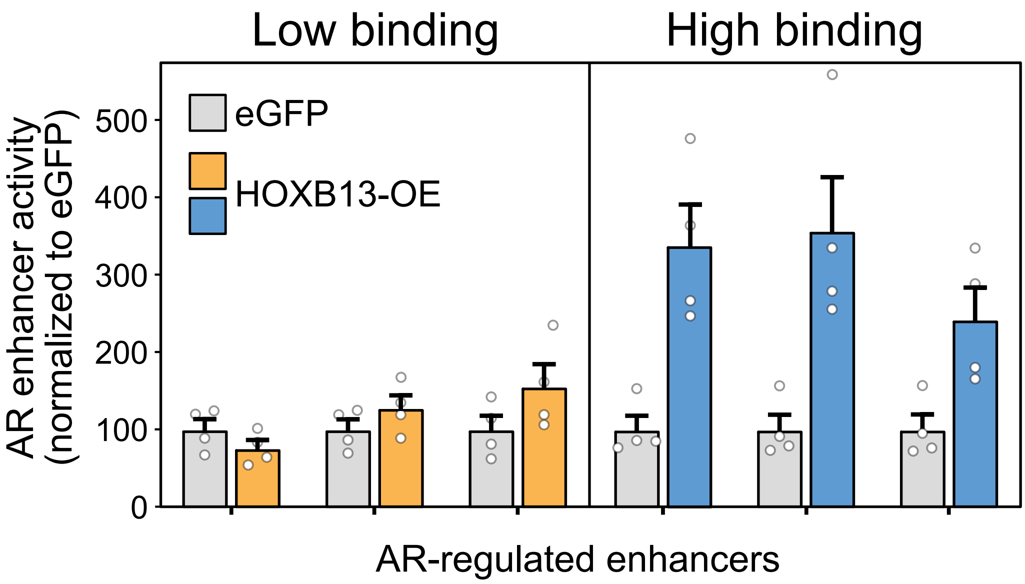


**Supplementary Figure 9:** Characterizing the impact of HOXB13 on AR-mediated enhancer activity. Validated AR enhancers (PMID: 33975627) were stratified into either HOXB131 “Low binding” and “High binding” regions based on LNCaP HOXB13 ChIP-seq (Supplementary Table 2). HEK293t cells were transfected with reporter plasmid, AR, and GFP or HOXB13 expression vectors. Enhancer activity was measured after 24h following 10nM DHT treatment and normalized to renilla transfection control. HOXB13 overexpression (OE) were compared to eGFP controls (n = 3 biological replicates; mean ± SEM).
